# Molecular identification of *Ageratum conyzoides* Linn (asteracea) leaf using DNA barcoding technique

**DOI:** 10.1101/2025.10.08.681161

**Authors:** Jacinta Enoh Apitikori-Owumi, Othuke Bensandy Odeghe, Marvellous Agbabune, Mubo Adeola Sonibare, Sayed Mohammed Firdous

## Abstract

**Background and Aims:** Reliable and accurate identification of medicinal plants is very pivotal for their safe use and handling in new drug development. This study is aimed at identifying the leaf of *A*. conyzoides Linn plant by using popular DNA barcode technique.

**Methods:** Genomic DNA was isolated from the plant sample followed its amplification through polymerase chain reaction convectional method. DNA barcoding specifically, Mat-K was used to identify and classify the plant sample. Consequently, plant sample was purified and sequenced. An evolutionary tree was constructed to assess *A. conyzoides* common ancestry.

**Results:** The plant sample showed a high degree of homology with known sequences in NCBI database after blasting analysis.

**Conclusions:** The findings indicated that DNA barcoding is a reliable and effective for medicinal plant identification and Maturase K (mat-K gene) is a reliable option for this application.

## INTRODUCTION

In the past few years, with the increase awareness in public health, herbal dietary supplement have gained growing acceptance among individuals owing to their therapeutic qualities. Researches showed that the global yearly sales of these herbal dietary supplements were forecast to increase and exceed US$140 billion as recorded in 2024. Nevertheless, the intricacy of species of plants and their financial values have led to unintentional and intentional adulteration of medicinal plants. It is noteworthy to say that herbal dietary supplements are governor by food law in USA and Europe while in some counties like Nigeria, they are taken as drugs (Li et al. 2024). Accurate species identification of the medicinal plants is a major aspect of quality control but accurate identification is dependent on morphological features such as shape, color, taste, texture, and scent, and accurate identification relies on the skills of the expert. It is important to note that morphological features are influenced by environmental and genetic factors, and therefore, expert identification of medicinal plants is subject to error (Hyder et al. 2024)

Several studies have shown that using only morphological characters for identification of plants results in errors. Therefore, the use of DNA-based technique would provide a definitive authentication to distinguish plant species. This new trend to rapidly identify and assess organisms, including plants, animals, and microorganisms, is based on the sequence of the DNA extracted from a tiny sample of the individual organism. It can also detect any new species that may be present, and is a potential tool for solving the error common to species identification, since this method often integrates similarity-based DNA barcoding and morphology to address taxonomic problems and any stage of plant growth can be used (Alade et al. 2023).

*Ageratum conyzoides* is a tropical species of plant in the family Asteraceae, commonly called goat weed, that originated in Central and South America and has spread to other tropical and subtropical areas, especially in Africa, Asia, and South America, growing in a variety of habitats, including grasslands, disturbed areas, roadsides, and agricultural fields (Joseph et al. 2024).

*Ageratum conyzoides* has traditionally been used for medicinal purposes and incorporated into the traditional medicine of diverse cultures; indigenous peoples from the native range of the plant have used the leaves, stems, and roots in fresh form or as a dried powder for a variety of health conditions, including wounds, fever, inflammation, respiratory infections, gastrointestinal disorders, and reproductive issues. *Ageratum conyzoides* is valued for its analgesic, anti-inflammatory, antimicrobial, antidiarrheal, and antipyretic effects and has been used as a decoction, infusion, poultice, or topical application by traditional healers and herbalists (Joseph *et al*. 2024).

## MATERIAL AND METHOD

### Plant collection and identification

Fresh plant sample **of** *Ageratum conyzoides* Linn was collected from the surrounding of Pharmacy, Delta State, Nigeria. Plant sample was identified and authenticate in the herbarium of the Department of Plant Biotechnology, University of Benin. A voucher specimen with a voucher number: UBH-A344 for *Ageratum conyzoides* Linn was deposited at the herbarium.

### Molecular identification of plant sample

Extraction of DNA of plant samples were done using a ZR Plant/seed Quick-DNA mini prep extraction kit supplied by Inqaba, South Africa. The process involved homogenizing 150 mg of respective plant samples. This was quickly followed by series of centrifugation of homogenized samples and various transfer steps were conducted out to purify the DNA. The isolated DNA was quantified using the Nanodrop 1000 spectrophotometer. SHV genes obtained from the isolates were amplified using the MATK-1RKIM-F: 5’ A CCCAGTCCATCTGGAAATCTTGGTT C-3’ and MATK-3FKIM-R: 5’-CGTACAGTACTTTTGTGTTTACGAG-3’ primers on an ABI 9700 Applied Biosystems thermal cycler at a final volume of 30 microliters for 35 cycles. The products were analyzed using 1% agarose gel electrophoresis at 200V for 15 minutes and visualized on a blue light imaging system for 900 bp product (Heckenhauer, 2016). Sequencing was done using the BigDye Terminator kit on a 3510 ABI sequencer by Inqaba Biotechnological, Pretoria South Africa. The sequencing was done at a final volume of 10ul, the components included 0.25 ul BigDye terminator v1.1/v3.1, 2.25ul of 5 x BigDye sequencing buffer, 10uM Primer PCR primer, and 2-10ng PCR template per 100bp. The sequencing conditions were as follows 32 cycles of 96°C for 10s, 55°C for 5s and 60°C for 4min (Diep, 2019). Phylogenetic analysis of the sequences was carried out using bioinformatics algorithm Trace edit, hit sequences downloaded from NCBI database using BLASTIN. Alignment of the sequence was done using ClustaIX. MEGA 6.0 was used for construction tree based on evolutionary history through Neigbor-Joining Method. The bootstrap consensus tree inferred from 500 replicates (Felsenstein, 1985) is taken to represent the evolutionary history of the taxa analysed. The evolutionary distances were computed using the Jukes-Cantor method (Jukes and Cantor, 1969).

## RESULTS

### Plants description

A description of *Ageratum conyzoides* plant used for molecular identification is shown in the table below.

**Table 1.** *Ageratum conyzoides* plant for isolation of DNA.

| Spice | Family | Part used | Medicinal properties |
| --- | --- | --- | --- |
| *Ageratum conyzoides* Linn. | Asteraceae | Leaf | Wound healing, inflammation |

### Agarose gel electrophoresis of MAT-k gene of *A. conyzoides* and *P. clematidea*

The purity of extracted DNA from *A. conyzoides* Figure 3) were determined by 1 % agarose gel electrophoresis. The sample showed a single band of three lanes, P, 1 and 2. 900 bp and 100 bp. Lanes 1-2 represent MAT-K gene bands (900 bp). Lane P represents the 100 bp Molecular ladder.

### Phylogenetic analysis of *Ageratum conyzoides* and *P. clematidea* plants samples

The obtained Mat-K sequence from the plants produced an exact match during the megablast search for highly similar sequences from the NCBI non-redundant nucleotide (nr/nt) database. The MatK of the plants showed a percentage similarity to other species at 100%. The evolutionary distances computed using the Jukes-Cantor method were in agreement with the phylogenetic placement of the MatK of the plants within the *Ageratum sp* revealed a closely relatedness to *Ageratum conyzoides* (Figure 2).

## 4.0 DISCUSSION

Accurate identification of medicinal plant species is a fundamental requirement for monitoring and conserving large-scale biodiversity. DNA barcoding is a rapid sequencing approach for identify a species that compares the sequence of an unknown specimen against barcodes in a reference database of known species based on similarity. Nowadays, four standard barcode regions (Mat-K, ITS, rbc and tmH-psbA) are routinely used for plant species identification through the DNA barcoding method. The Mat-K region is an effective and reliable DNA barcoding for identifying medicinal plants of varied geographical origins.

This study is aimed at identifying *A*. conyzoides plant using popular DNA barcode technique. The 900 bp bands observed for the DNA extracted from both plants suggested that the polymerase chain reaction (PCR) was successful in amplifiying of the Mat-K gene, the targeted gene, consequently shows uncontaminated DNA (Alade *et al*., 2023). The size of the band (900 bp) corresponds to the expected size of the target gene suggesting specificity in the PCR reaction. The small size of the brand (100 bp) might indicate the amplification of a short fragment or specific region within the gene as a result of contamination of the chloroplast DNA or degraded DNA. The sequences obtained by direct sequencing of the PCR product was of high quality, with simple-to-score evenly spaced peaks in the sequence traces. MatK may be a starting point for quality control and assurance of plant materials used in research, manufacturing, customs, and forensics (Abdelsalam et al. 2022). The Maturase K (matK) gene of plant chloroplast is a strong marker in studying plant molecular systematics, evolution, and genetic polymorphism and is considered as the standard DNA barcode for plants. The gene has an important role in the phylogenetic reconstruction of terrestrial plant (Rani, 2023). The Maturase K (mat-k) gene has recently become popular in plant molecular systematics and evolution because of the genes rapid evolution at nucleotide and corresponding amino acid levels (Barthet et al. 2020). The degree to which a plant species resembles one that has already been identified in the database is known as maximum identity. Accuracy of identification is defined as a maximum similarity of at least 95 percent (Alade *et al*., 2023). One hundred percent (100%) maximum identity was found for *A. Conyzoides* (Figure 1) following the blasting the samples against NCBI database. The 100% identity of *A. conyzoides* match suggests that the query sequence is from same species as the target (subject) sequence which can be useful for identifying plant species. High percentage identity matches can indicate conserved species which can provide insights into the revolutionary relationships between organisms. Furthermore, the high identity matches can be useful in various applications such as phylogenetic analysis, gene cloning or diagnostic testing. The research on medicinal plants genomes has improved rapidly through DNA barcoding and high-throughput sequencing and their contributions to enhancing the accuracy and reliability of plant identification and limitation by their huge genome size and high repetitive sequence has been overcome (Cheng *et al*., 2021, Wang *et al*., 2024). Indisputably, genomic regions differ significantly in their potential phylogenetic explanatoriness and their inputs in solving a taxonomic group over a designated period (Udensi *et al*., 2017)

**Figure 1.**
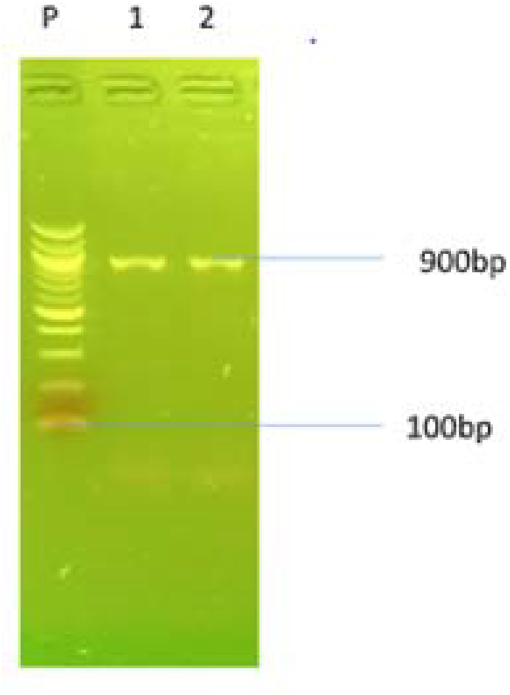
Agarose gel electrophoresis of MAT-k gene of *Ageratum conyzoides*. Lanes 1-2 represent MAT-K gene bands (900 bp). Lane P represents the 100 bp Molecular ladder.

**Figure 2.**
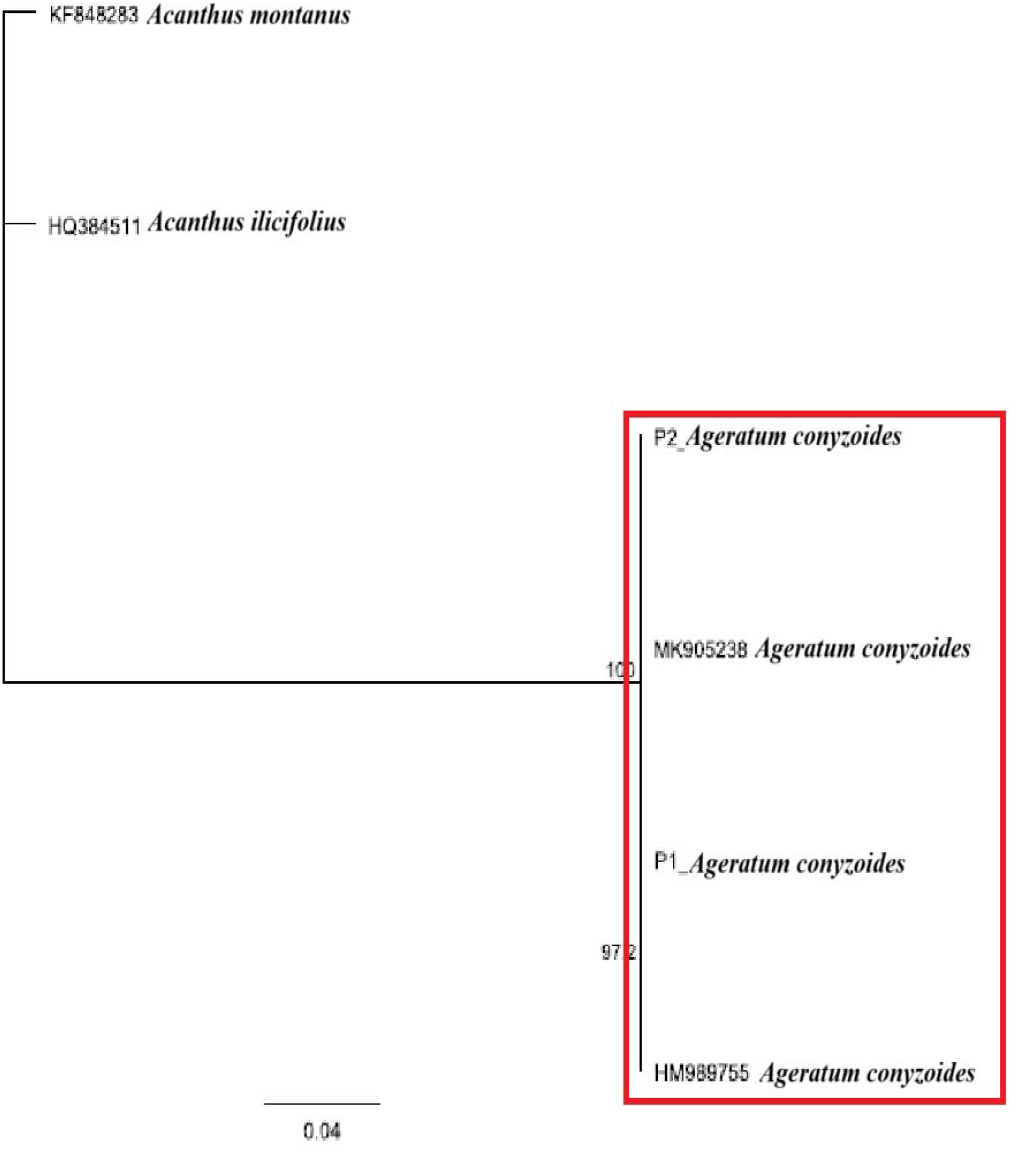
Phylogenetic tree of *Ageratum conyzoides*.

## CONCLUSION

The results from this study successfully showed that DNA barcoding is a robust technique and reliable application for identifying local medicinal plant. Likewise, *A. conyzoides* identification in this study will be helpful in the development of new experiments with other medicinal plants in Delta State.

## Informed Consent

Informed consent was obtained from authors

## Authors’ contributions

A.J.E, M.A.S, O.B.O, M.A: Concept/Design, A.J.E: Drafting Manuscript: A,J.E/S.M.F: Results Interpretation

